# Targeted Sortase A Inhibition by Novel Peptidomimetic Antivirulents against Staphylococcal Infections

**DOI:** 10.1101/2025.01.14.632915

**Authors:** Jordi C. J. Hintzen, Shadi Rahimi, Daniel Tietze, Jian Zhang, Ivan Mijakovic, Alesia A. Tietze

## Abstract

Antibiotic resistance is a critical public health issue, causing resistant bacterial strains to be increasingly difficult to control. Antivirulence therapies, which target bacterial virulence factors rather than kill bacteria, present a promising approach. Sortase enzymes, particularly SrtA, are crucial for Gram-positive bacterial virulence by anchoring surface proteins essential for bacterial adhesion and biofilm formation to the bacterial outer cell wall. This study evaluates the selectivity of the peptidomimetic inhibitor BzLPRDSar towards various Gram-positive bacteria. The BzLPRDSar significantly inhibited biofilm formation in multidrug-resistant *S. aureus* and *S. epidermidis*. Conversely, it showed variable and generally lower selectivity to Gram-positive species such as *E. faecalis, B. cereus* and *S. agalactiae*. The selectivity towards Staphylococcus species is attributed to conserved structural elements in the SrtA enzyme, particularly the β7/β8 loop region with a key tryptophan, likely facilitating strong binding interactions with the inhibitor.

**Importance:** This study addresses the pressing issue of antibiotic resistance by exploring antivirulence therapy as an innovative alternative to conventional antibiotics, focusing on inhibiting bacterial virulence rather than bacterial growth. By evaluating the selectivity of the peptidomimetic inhibitor BzLPRDSar against various Gram-positive bacteria, the study highlights its potent selectivity in inhibiting biofilm formation in multidrug-resistant *S. aureus* and *S. epidermidis*. The findings underscore the potential of targeting conserved structural elements in bacterial Sortase enzymes, particularly in Staphylococcal species, to develop more selective and effective antivirulence therapies.

## Introduction

Antibiotic resistance has become one of the most pressing public health challenges of the 21^st^ century. The overuse and misuse of antibiotics in both human medicine and agriculture have accelerated the evolution of resistant bacterial strains. This resistance complicates the treatment of common infectious diseases and increases the risk of spreading resistant pathogens.^1^ For example, methicillin-resistant *Staphylococcus aureus* (MRSA) is responsible for severe infections that are difficult to treat and lead to high morbidity and mortality rates.^2^ Similarly, multidrug-resistant *Enterococcus faecalis* and other Gram-positive bacteria pose significant treatment challenges, often requiring the use of last resort antibiotics, further accelerating the development of resistance against these treatments.^3^ The development of new antibiotics has significantly slowed down over the past few decades due to high cost and lengthy process of antibiotic research and development, coupled with lower financial incentives for pharmaceutical companies.^4, 5^ Moreover, the rapid emergence of resistance to new antibiotics further disincentivizes investment in antibiotic research. Given these critical issues, alternative strategies to combat bacterial infections are urgently needed. One promising approach is the development of antivirulence therapies. Unlike traditional antibiotics, which aim to kill bacteria, antivirulence therapies target the virulence factors bacteria use to establish infections and cause disease.^6-8^ By disarming the bacteria rather than killing them, these therapies exert less selective pressure for the development of resistance. This approach includes targeting specific bacterial processes such as toxin production, adhesion to host tissue and biofilm formation.^9, 10^

Among the virulence factors, sortase enzymes, found in all Gram-positive bacteria, are among the most well-studied. Sortases are a class of cysteine proteases responsible for anchoring surface proteins to the outer bacterial cell wall.^11^ Within the family of sortase enzymes found in different bacterial species sortases A to F were described and differ in their primary sequence.^12^ Furthermore, sortase subtypes differ in their target proteins and have distinct biological roles. Within the target proteins, sortases recognize a short motif consisting of specific amino acids, where the enzyme can cleave and react with the substrate protein.^13^ These recognition motifs further define the subtype of a specific enzyme which leads to the distinct roles of the subtypes within a bacterial species.

Arguably the most important among this class of enzymes is sortase A (SrtA), which recognizes the LPxTG motif and anchors its target proteins to a Lipid II molecule (Fig. 1A).^13^ The target proteins of SrtA are involved in bacterial adhesion to the host tissue and are defined by the umbrella term microbial surface components recognizing adhesive matrix molecules (MSCRAMMs). These MSCRAMMs adhere to the hosts’ extracellular matrix proteins, such as fibrinogen and fibronectin, and are, therefore, crucial for the bacteria to invade their host.^14^ Additionally, SrtA is also required for proper adhesion of the bacteria amongst each other and plays an important role in the formation of pathogenic biofilms of many Gram-positive species (Fig. 1A).^15^ These roles in bacterial physiology make SrtA a prime target for antivirulent therapy, as inhibition of this enzyme would lead to prevention of host tissue adherence as well as disruption of biofilm formation, leading to greatly reduced pathogenicity.

**Fig. 1:**
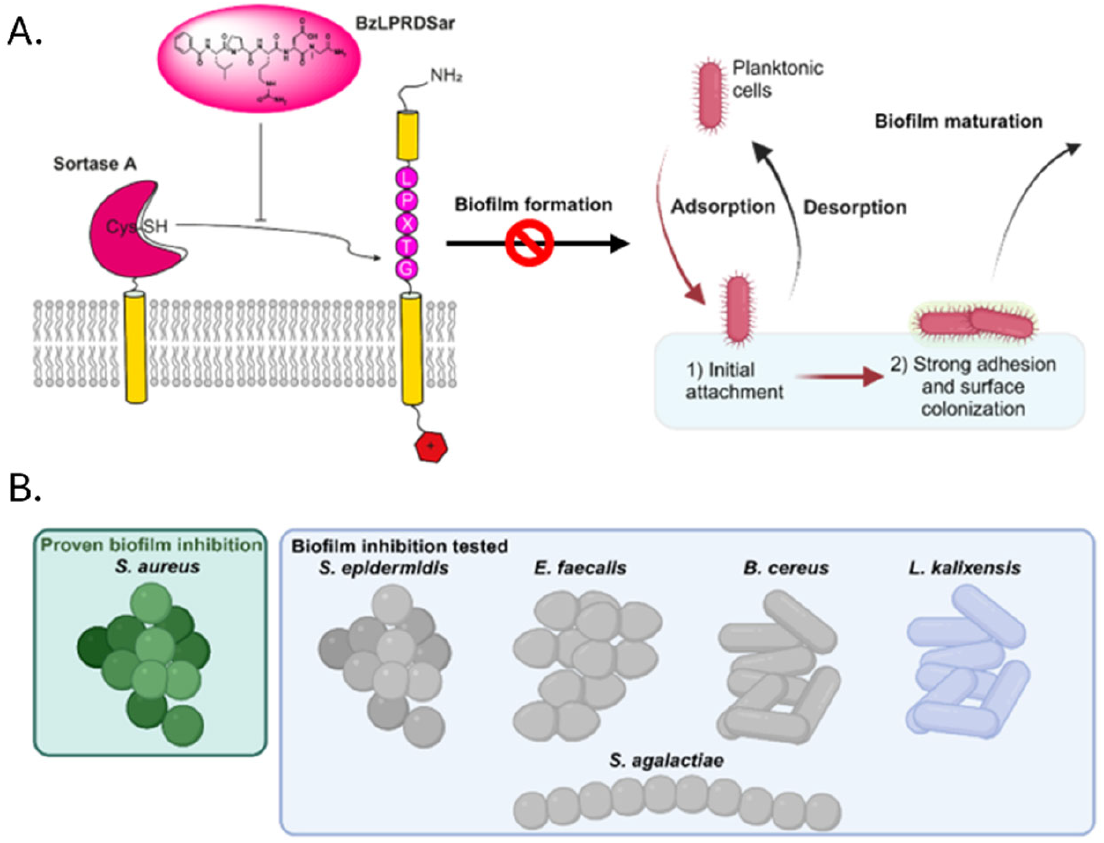
**A)** Mechanism of action of peptidomimetic SrtA inhibitors to prevent biofilm formation; **B)** Overview of Gram-positive strains that have been tested for biofilm inhibition by BzLPRDSar in this study.

Diseases caused by Gram-positive bacteria are often severe and highlight the critical need for new therapeutic strategies. For example, MRSA is notorious for its resistance to multiple antibiotics, making it a significant threat in both healthcare and community settings.^2^ Especially in infections like these, antivirulence could provide a crucial new treatment option. One disease which has been directly associated with SrtA is septic arthritis.^16^ Septic arthritis is a severe infection of the joints that can lead to permanent damage if not treated effectively.^17^ Here, SrtA inhibitors could serve as a first in line treatment option to combat septic arthritis. Biofilm formation is a common strategy employed by bacteria to protect themselves from the host immune system and antibiotics.^18^ Biofilms are complex communities of bacteria that adhere to surfaces and are embedded in a self-produced extracellular matrix. This matrix shields the bacteria from environmental stresses, making infections difficult to eradicate. Therefore, targeting biofilm formation, for example by inhibiting SrtA, is a critical aspect of developing effective antivirulence therapies.

SrtA inhibitors have been an active research field, with numerous studies exploring their potential to reduce bacterial virulence and biofilm formation.^19-22^ Previously, we reported on the synthesis and evaluation of novel peptidomimetic substrate-derived SrtA inhibitors (Fig. 1A).^23^ These compounds were active against expressed *S. aureus* SrtA with low micromolar IC_50_ values *in vitro*. Importantly, three compounds, BzLPRDSar, FLPRDA and BzLPRDF inhibited the growth of *S. aureus*. It was observed that the bacteria could still survive, but the plateau of the growth curve was significantly lower, suggesting a reduced ability to adhere to each other. BzLPRDSar, the compound with the largest effect on bacterial growth, was also tested for its ability to inhibit the biofilm formation of *S. aureus* and was able to eliminate 95% of the formed biofilms at 128 μg/ml.

Encouraged by the results when targeting *S. aureus*, we have now explored the potential of our lead peptidomimetic SrtA inhibitor, BzLPRDSar, against a variety of pathogenic and multidrug resistant Gram-positive bacteria (Fig. 1B). In our panel, we included a multidrug resistant strain of *S. aureus* to verify the efficacy of our compounds in comparison to a wild-type *S. aureus* strain as was used earlier. Another multidrug resistant bacterial species that was included is *E. faecalis. E. faecalis* is known to cause life threatening infections such as endocarditis, sepsis, urinary tract infections and meningitis. Particularly, *E. faecalis* forms persistent antibiotically resistant biofilms, making it an excellent target for SrtA inhibition.^24, 25^ *Bacillus cereus* can cause foodborne illness due to its spore-forming nature, furthermore biofilms formed by *B. cereus* pose a large challenge in the food production industry as it grows readily on air-liquid interfaces and hard surfaces.^26^ Closely related to *B. cereus* is the species *B. anthracis*, which is well known as the anthrax bacteria and classified as a major pathogen within the *Bacillus* genus.^27^ The next species we included in the panel, *Lactobacillus kalixensis*, is known to form biofilms in the vaginal and gut microbiota, where it can become pathogenic.^28^ Importantly, *L. kalixensis*, as with other lactobacilli, grows under anaerobic conditions. As a representative of the *Streptococcus* genus, *Streptococcus agalactiae* was included, which is the most common human pathogen of the streptococci. It can cause severe invasive infections in elderly, new-borns and immunocompromised patients.^29^ *S. agalactiae* is also a common bovine pathogen as it is the main cause of bovine mastitis, making treatment of this bacterium very relevant for veterinary medicine as well.^30^ Finally, *Staphylococcus epidermidis*, a species found in the normal skin and sometimes mucosal microbiota was included. In some cases, *S. epidermidis* can cause dangerous infections and is notoriously hard to treat.^31^

Comparing these bacterial species gives us a wide variety of distinctly different pathogenic Gram-positive bacteria ranging from several major families, providing insight into the potential of our peptidomimetic SrtA inhibitors as a pan-SrtA inhibitor, which could be applied to treat widely different infections. Furthermore, these pathogens are involved in many different disease phenotypes in humans but also stretch to veterinary medicine and the food industry, conceivably increasing the scope of the potential of these inhibitors. Therefore, we have investigated the effect of our previously discovered lead compound, BzLPRDSar, on the growth and biofilm formation of these bacteria.^23^

## Results and discussion

From our initial panel, several compounds were able to affect the growth of *S. aureus* bacteria, however, BzLPRDSar stood out among these as the compound with the largest effect on the growth profile of *S. aureus* and additionally, was able to efficiently disrupt biofilm formation of this species at concentrations as low as 32 μg/mL, making it the compound of choice for testing towards other bacterial species.^23^ It was synthesized and purified according to previously reported procedures and subsequently used in the bacterial assays (Fig. S1). For the growth profiling assays, the different bacterial species were grown in Tryptic Soy Broth (TSB), transferred to a 96-well plate and subsequently incubated with varying concentrations of BzLPRDSar, at 2, 8, 32 and 128 μg/mL. To allow the multidrug resistant species to grow optimally, *S. aureus* and *E. faecalis* and *K. pneumoniae* were grown and incubated in Mueller Hinton Broth (MHB), while *L. kalixensis* was grown and incubated under anaerobic conditions in TSB in an atmosphere consisting of 5% CO_2_ and 6% O_2_. As a negative control, the Gram-negative bacterial species *Klebsiella pneumoniae* was included in the panel, which lacks SrtA all together and is, therefore, not expected to be affected by our peptidomimetic inhibitor. Overall, the same trend as reported previously was observed, where the bacteria grew in the first 6 to 12 hours undisturbed (Fig. 2) and only after reaching their respective plateaus, a difference in their growth profile could be observed. Interestingly, different bacterial species showed vastly different growth profiles, but consistently, the plateau was reached after 48 hours of incubation and was used as a measure to define their growth decrease when treated with BzLPRDSar. Comparatively, the MDR strain of *S. aureus* was affected to a lesser extent than the previously used wild type strain (Fig. 2A).^23^ Whereas BzLPDRSar displayed a 36% decrease in growth at 128 μg/mL for the wildtype strain, only a 10% decrease could be observed for the MDR strain, suggesting its more persistent nature (Table 1). However, other bacterial species were efficiently inhibited in their growth at this concentration, with multidrug resistant *E. faecalis, B. cereus* and *S. epidermidis* showing moderate growth inhibition at around 20% (Table 1, Fig. 2B-D). *S. agalactiae* proved to be the species least efficiently affected amongst the tested Gram-positive species, showing only an 8.9% decrease at 128 μg/mL (Table 1, Fig. 2E). Fortunately, the Gram-negative *K. pneumoniae* was not significantly inhibited, showing a negligible decrease in growth at even the highest inhibitor concentration present, further confirming the presence of SrtA in Gram-positive species is needed for efficient growth inhibition (Table 1, Fig. S2). Finally, *L. kalixensis* was the species that was most affected by our peptidomimetic inhibitor, showing a 42% decrease in growth at 128 μg/mL, outperforming the originally tested *S. aureus* strain (Table 1, Fig. 2F). Taken together, these results show that Gram-positive bacteria are indeed specifically targeted by BzLPRDSar at varying degrees differing from species to species. However, as all these species grow pathogenic biofilms that can cause major threats in health, the ability of BzLPRDSar to disrupt these biofilms was assayed for all Gram-positive species.

**Table 1.** Growth profiling overview.

| Species | Decrease in growth 32 $\mu\text{g/ml}$ | Decrease in growth 128 $\mu\text{g/ml}$ |
| --- | --- | --- |
| *Staphylococcus aureus* MDR | 6% ± 1% | 10% ± 2% |
| *Staphylococcus epidermidis* | 16% ± 4% | 22% ± 1% |
| *Enterococcus faecalis* MDR | 6% ± 5% | 23% ± 3% |
| *Bacillus cereus* | 4% ± 7% | 20% ± 7% |
| *Streptococcus agalactiae* | 5% ± 2% | 9% ± 1% |
| *Lactobacillus kalixensis* | 17% ± 4% | 42% ± 3% |
| *Klebsiella pneumoniae* MDR | 2% ± 1% | 2% ± 1% |

**Fig. 2:**
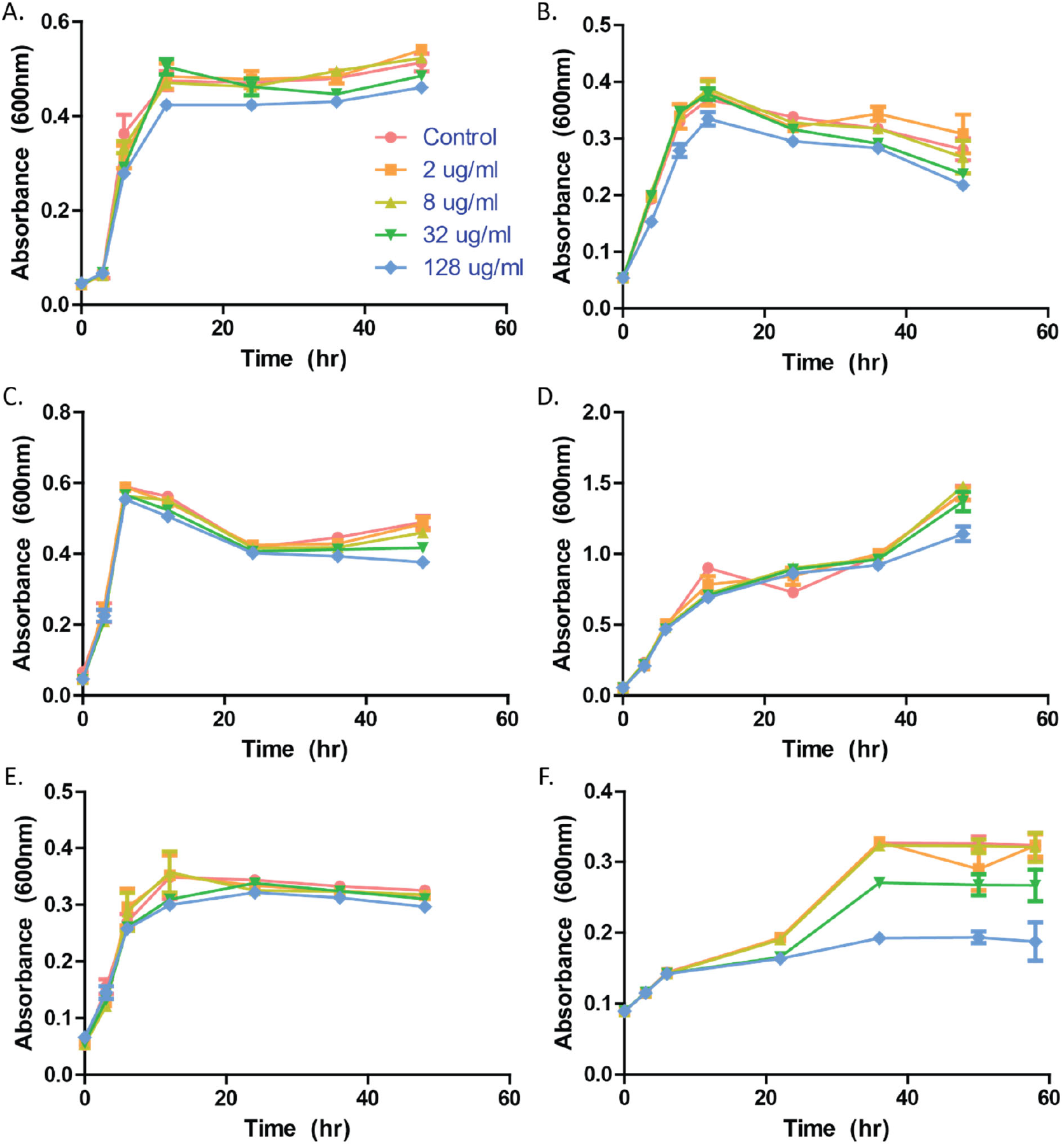
Growth profiling curves of bacteria incubated in presence of varying concentrations of BzLPRDSar. The absorbance was measured at 600 nm as a measure of OD for **A)** *S. aureus* MDR; **B)** *S. epidermidis;* **C)** *E. faecalis* MDR; **D)** *B. cereus*; **E)** *S. agalactiae*; **F)** *L. kalixensis*. All samples were measured in triplicate, error bars are reported as standard error (±SE).

Bacterial biofilms were generated as previously described by incubating varying concentrations of BzLPRDSar in Brain Heart Infusion Broth on coagulase coated 96-well plates.^23^ After an overnight incubation with the peptidomimetic inhibitors, bacterial biofilms were stained with crystal violet (CV) and solubilized in 90% ethanol to quantify the remaining amount of biofilm by measuring the absorbance at 595 nm. As was reported before, *S. aureus* biofilms were efficiently eradicated at concentrations of 32 and 128 μg/mL of BzLPRDSar, in line with previously reported values, with 34% and 26%, respectively, of remaining biofilm (Fig. 3, Table 2). In line with growth inhibition, the MDR *S. aureus* strain used in these experiments proved to be slightly more resistant to treatment with the peptidomimetic than its wildtype counterpart. Surprisingly, an efficient disruption of biofilm formation was also observed for *S. epidermidis* at these inhibitor concentrations, with 48% and 36% of biofilm remaining, respectively (Table 2, Fig. 3). Seemingly, BzLPRDSar only had a moderate effect on the growth profile of *S. epidermidis* but was able to disrupt the formation of biofilms quite efficiently in this species. Among the other included bacterial species, biofilm inhibition proved to be less efficient than for the two *Staphylococcus* species tested, with around 60 to 70% of biofilm remaining for *E. faecalis, B. cereus* and *L. kalixensis* at 128 μg/mL and even 90% for *S. agalactiae* at this concentration (Table 2, Fig. 3). While *L. kalixensis* showed promising results when examining its growth profile, the biofilms formed by this bacterial species proved to be resistant towards treatment with our peptidomimetic inhibitor (Table 2, Fig. 3). All in all, it can be observed that a certain degree of selectivity is displayed in the biofilm inhibition experiments, where BzLPRDSar was efficient in disrupting biofilms of the genus of *Staphylococcus*, but did not show a considerable effect on biofilms of other species, providing an opportunity to selectively use these compounds to target *Staphylococci* infections.

**Table 2.** Biofilm inhibition overview.

| Species | Biofilm remaining 32 µg/ml | Biofilm remaining 128 µg/ml |
| --- | --- | --- |
| *Staphylococcus aureus* MDR | 34% | 26% |
| *Staphylococcus epidermidis* | 48% | 36% |
| *Enterococcus faecalis* MDR | 74% | 66% |
| *Bacillus cereus* | 79% | 61% |
| *Streptococcus agalactiae* | 93% | 90% |
| *Lactobacillus kalixensis* | 69% | 65% |

**Fig. 3:**
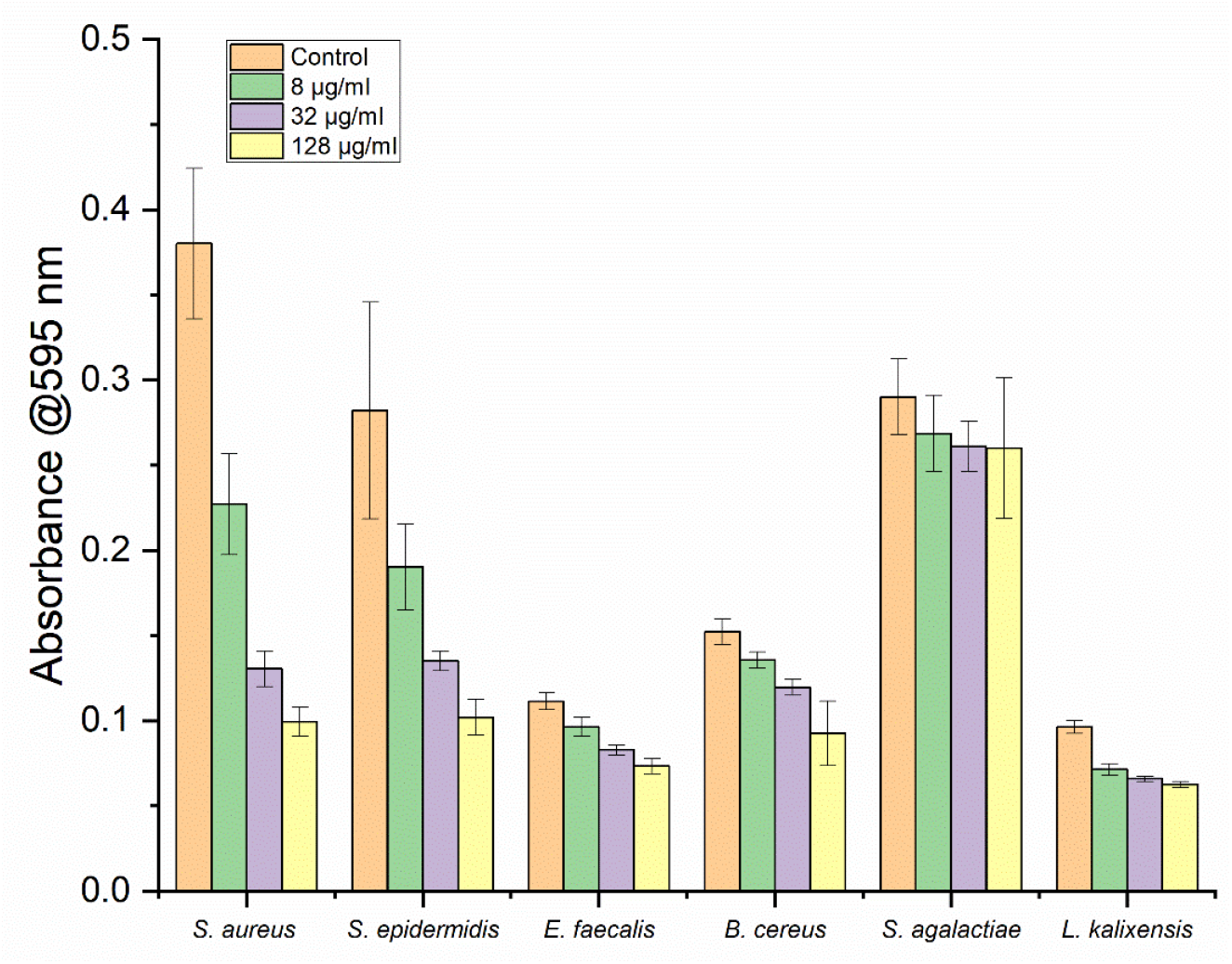
Absorbance measurements of bacterial biofilms stained with Crystal Violet which were incubated with varying concentrations of BzLPRDSar. Absorbance measured at 595 nm; samples were taken in triplicate, error bars reported as standard error (±SE).

To elucidate the molecular basis for the observed specificity of BzLPRDSar towards staphylococcus species, we conducted a detailed sequence alignment of the SrtA enzymes from the various bacterial species included in our study using Jalview and ClustalΩ.^19, 32, 33^ Strikingly, several key structural elements are conserved among the aligned staphylococcus species but vary greatly among the other evaluated species. The β2/H1 loop region which is known to be involved in substrate binding consists of the neutrally charged PATP sequence in *S. aureus* and *S. epidermidis*, while in many other species negatively or positively charged amino acids appear.^34, 35^ Similarly, the β6/β7, also involved in substrate binding, is most conserved between the staphylococci and varies greatly among the other species.^36, 37^ Perhaps most strikingly is the β7/β8 loop region, which is extended in a similar way only in the staphylococcal SrtA proteins, while for the other species this region is significantly shorter. It was shown earlier that the β7/β8 loop region has an influence on the binding of peptidomimetic inhibitors and therefore is suspected to play a role in the selectivity of BzLPRDSar as well.^37^

The alignment also revealed a notable difference in the amino acid sequences in the β7/β8 loop, particularly highlighting a single tryptophan residue at position 194 for *S. aureus* (W194) or position 191 for *S. epidermidis* (W191) in the elongated β7/β8 loop (Fig. 4A). The tryptophan residue present here in *Staphylococcus* species is absent in the SrtA sequences of the other Gram-positive bacteria tested in our panel, such as *E. faecalis, B*. cereus and *S. agalactiae*. Our computational analysis conducted earlier suggested that the aromatic side chain of the tryptophan residue at this position forms some key interactions with the Sar residue of the BzLPRDSar molecule (Fig. 4B).^23^ This interaction likely stabilizes the binding of the inhibitor to the active site of the enzyme, enhancing its inhibitory potency. In the absence of this tryptophan residue in other bacterial species, such stabilizing interaction are not possible, which could explain the reduced efficacy of BzLPRDSar against these non-staphylococcal SrtA enzymes (Fig. 4A, B). Also, the physicochemical properties and thus the overall shape of the active site varies slightly among the different species as can be concluded from the sequence alignment and Fig. 4 C,D. As stated earlier, the flexible β7/β8 loop has a strong influence on the shape and properties of the binding site and seems to play a key role in inhibitor binding similarly as it is a major player to alter substrate specificity among sortase enzymes.^40^ An exception is *L. kalixensis*, which showed promising results in the growth inhibition assays, but only moderate biofilm inhibition. While *L. kalixensis* does possess a tryptophan residue in the active site, but in a slightly different location. Moreover, the surrounding defining loop regions (Fig. 4A) are vastly different from SrtA from *S. aureus* and *S. epidermidis*, providing a reasonable explanation for the difference in effectivity of BzLPRDSar.

**Fig. 4:**
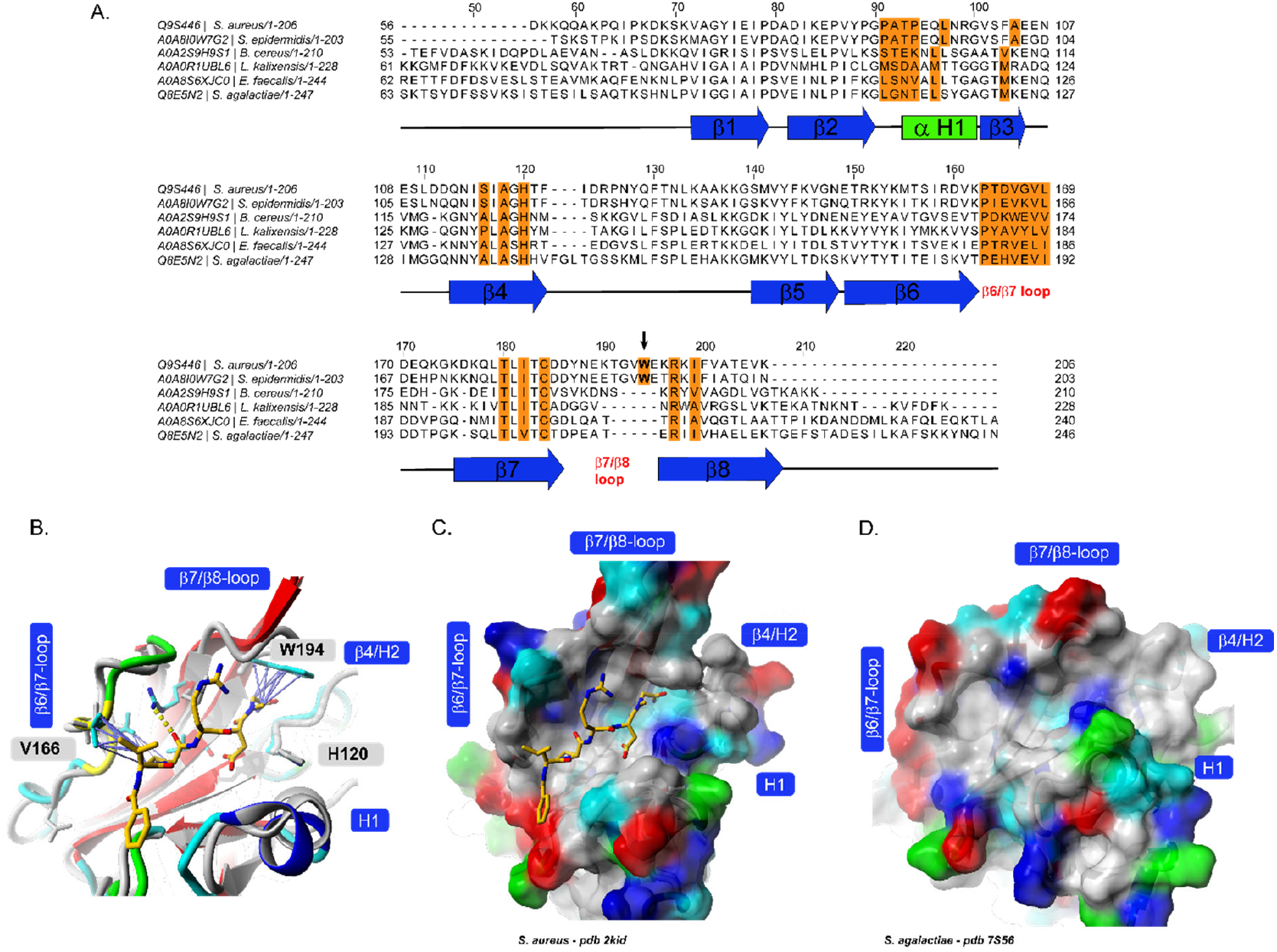
**A)** Sequence alignment of the SrtA enzymes from the studied bacterial species. Highlighted in orange are residues in and around the substrate binding site, while secondary structure elements are indicated with lines and arrows underneath the sequence alignment. W194 as a potential key residue for inhibitor binding affinity is indicated with an arrow. **B)** structural alignment of *S. agalactiae* (grey) and *S. aureus* SrtA bound to BzLPRDSar (coloured) highlighting some key Van-d-Waals interaction (purple arrows) to W194 and V166 (protein structures shown in ribbon style, ligand and some interacting protein side chains shown in stick representation, colour scheme: ligand carbon – orange, protein side chain carbon – cyan, nitrogen – blue, oxygen - red). **C, D)** surface representation of *S. aureus* SrtA with bound inhibitor (same colouring scheme as in B) and *S. agalactiae* SrtA (surface coloured according to the physicochemical properties of the amino acids, colour scheme: polar – cyan, green, unpolar – grey, red – acidic, blue - basic). Structure views created with Yasara.^38, 39^

The highly similar loop regions and unique presence of this tryptophan in staphylococcus species provides a structural rationale for the selective inhibition observed in our growth and biofilm assays. The strong interaction between BzLPRDSar and the W194 residue effectively aids disruption of the SrtA function in *S. aureus* and *S. epidermidis*, leading to the observed significant reductions in bacterial growth and biofilm formation. In contrast, the lack of this residue in other species in combination with critical differences in the loop regions results in a less effective binding of the inhibitor, thereby reducing its impact on their growth and biofilm development.

Finally, we performed Scanning Electron Microscopy (SEM) imaging for *S. aureus* and *S. epidermidis* biofilms to observe potential structural changes in the bacterial cellular membrane and the interbacterial association within the biofilm.^41^ Earlier, confocal microscopy could confirm some differences in overall colony formation and morphology of *S. aureus* treated with peptidomimetic inhibitors.^23^ Staphylococcus biofilms were grown directly on SEM glass cover slides and incubated with BzLPRDSar at a concentration of 128 μg/ml, a concentration that could eliminate a major part of the biofilms when grown on a coagulase substrate (Table 2). After 18 hours of incubation with the peptidomimetic compound, the bacteria were fixated using glutaraldehyde and subsequently incubated with increasing concentrations of ethanol, ranging from 40% to 100%. After the final ethanol step, the samples were dried and coated with a 5 nm gold layer and directly measured.

SEM measurements revealed notable differences in the structural integrity and interbacterial associations within the biofilms of *S. aureus* and particularly of *S. epidermidis* when treated with BzLPRDSar. For both species, the untreated biofilms displayed a typical multi-layered structure with tightly associated bacterial cells (Fig. 5, Fig. S3).^42^ The cellular membranes of the treated *S. aureus* appeared slightly smoother compared to the untreated control, suggesting potential membrane remodelling or stress responses induced by the inhibitor (Fig. 5A). For *S. epidermidis*, the treated biofilm structure was markedly disrupted. The cells appeared more globular and less interconnected, indicating a significant reduction in interbacterial adhesion and biofilm integrity (Fig. 5B). These structural changes confirm the biofilm inhibition data, highlighting the efficacy of BzLPRDSar in disrupting biofilm formation in staphylococcus species.

**Fig. 5:**
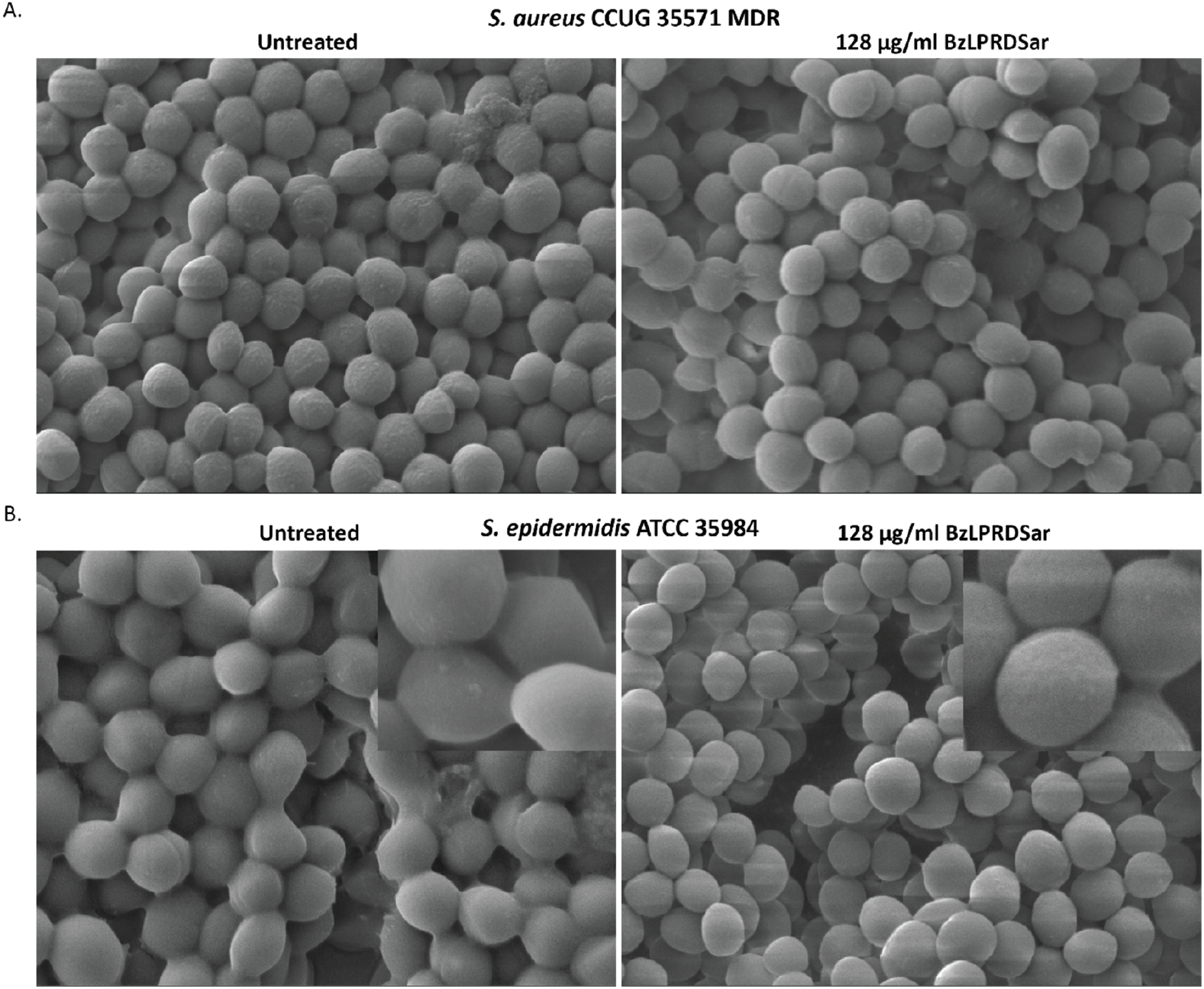
SEM images illustrating structural changes in biofilms of *S. aureus* and *S. epidermidis* incubated with 128 μg/ml of BzLPRDSar for 18 hours. Magnification 15,000x, with insets at 50,000x. **A)** SEM images of *S. aureus* biofilms, untreated control (left) and treated biofilm (right), **B)** SEM images of *S. epidermidis* biofilms, untreated control (left) and treated biofilm (right).

In this study, we evaluated the selectivity of our lead peptidomimetic Sortase A (SrtA) inhibitor, BzLPRDSar,^23^ in inhibiting the growth and biofilm formation of a variety of pathogenic Gram-positive bacterial species. Our findings indicate that BzLPRDSar is highly effective in inhibiting both the growth and biofilm formation of *S. aureus*, including the evaluated MDR strain, as well as *S. epidermidis*. BzLPRDSar disrupted biofilm formation to a notable extent, with only 26% and 36% of biofilm remaining for *S. aureus* and *S. epidermidis*, respectively, at the highest concentration of 128 μg/ml tested. The selective inhibition can be attributed to the unique structural features of the peptidomimetic compound. Sequence alignment revealed that staphylococcal SrtA enzymes possess a tryptophan residue near the active site, which appears to be specifically targeted by the C-terminal sarcosine (Sar) residue of BzLPRDSar. This molecular interaction likely enhances the binding affinity and inhibitory potency of BzLPRDSar against staphylococcal SrtA, explaining its preferential activity. The identification of the W194 residue (or W191 in *S. epidermidis*) highlights our rational design strategy for inhibitor design in developing targeted antimicrobial therapies. This approach not only enhances the selectivity of the inhibitors against specific pathogens but also can minimize off-target effects. These findings provide a compelling explanation for the selective activity of BzLPRDSar and pave the way for further optimization and development of targeted SrtA inhibitors. Additionally, SEM analysis of *S. epidermidis* treated with BzLPRDSar revealed that the association between bacteria was significantly disrupted. This morphological evidence supports the hypothesis that BzLPRDSar impairs the adhesion processes mediated by MSCRAMMs, which are anchored to the cell wall by SrtA. By inhibiting SrtA, BzLPRDSar prevents the proper attachment of these surface proteins, leading to a failure in biofilm formation and inter-bacterial cohesion.

All in all, our results demonstrated that BzLPRDSar is selective towards *Staphylococcus* species, which could provide an opportunity for developing targeted therapies that can effectively combat staphylococcal infections without broadly disrupting the microbiome. This selective action could prevent the common side-effects associated with broad-spectrum antibiotics, such as dysbiosis and disruption of beneficial microbial communities. However, *in vivo* studies are essential to validate the therapeutic potential of these compounds in animal models of staphylococcal infections. Encouragingly, a related peptidomimetic SrtA inhibitor, LPRDA, was recently shown to be efficient *in vivo* in the treatment of peritoneal infections of *S. aureus* in mice, highlighting the potential of this class of compounds even further.^43^ The development of these targeted antivirulence therapies has the potential to revolutionize the treatment of bacterial infections, offering a new strategy to combat antimicrobial resistance.

## Methods

### Peptide synthesis and purification

Peptidomimetic inhibitor BzLPRDSar was synthesized according to previously reported procedures.^23^ Briefly, the peptide was chain assembled using Rink amide resin manually. Amino acid couplings were carried out with the molar ratio of (4):(4):(8) of (Fmoc-protected amino acid):(HATU):(diisopropylethylamine) at room temperature for 60 minutes and deprotection was achieved in 20% piperidine in DMF for 30 minutes at room temperature. N-terminal benzoylation was achieved with benzoyl chloride (10 eq.) and triethylamine (10 eq.) in DMF for 2 hours at room temperature. The peptide proceeded to standard cleavage from resin using a mixture of trifluoroacetic acid (TFA, 95%), H_2_O (2.5%) and triisopropylsilane (2.5%) for 3 hours at room temperature. TFA was removed using N_2_ and the resultant residue suspended in ice-cold diethyl ether. The mixture was then centrifuged (5 minutes, 4600 RPM) after which the supernatant was decanted into the waste. The remaining solid was washed twice by ice-cold diethyl ether and subjected to purifications. For the purification and characterization of peptides, two eluent systems were used. Mobile phase A was 0.1% TFA in MQ-H_2_O, mobile phase B was 0.1% TFA in acetonitrile (ACN). The crude peptide was dissolved in a mixture of ACN in H_2_O and purified by semi-preparative HPLC using a Waters 600 system equipped with a C18 column (Multokrom 100-5 C18, 5 μm particle size, 100 Å pore size, 250 × 20mm) and a gradient of mobile phase A and mobile phase B from 20% B to 60% B over 45 minutes at 8 mL/min. Detection of chromatographic peaks was analysed at 214 and 254 nm. Analytical RP-HPLC was carried out on a Waters XC e2695 system employing a Waters PDA 2998 diode array detector equipped with a ISAspher 100-3 C18 (C18, 3.0 μm particle size, 100 Å pore size, 50 × 4.6mm) at a flow rate of 2 mL/min using a gradient of mobile phase A and mobile phase B from 20% B to 60% B over 10 minutes. The molecular weight of the purified peptide was confirmed by ESI mass on a Waters Synapt G2-Si ESI mass spectrometer equipped with a Waters Acquity UPLC systems using a Xela C18 column (C18, 1.7 μm particle size, 80 Å pore size, 50 × 3.0mm).

### Growth profiling assays

For the growth profiling experiments, *S. aureus* strain CCUG 35571, *E. faecalis* strain CCUG 34289 and *K. pneumoniae* strain CCUG 37387 were cultured overnight in sterile MHB medium, while *B. cereus* strain UW85, *S. agalactiae* strain CCUG 4208T and *S. epidermidis* strain ATCC 35984 were cultured overnight in sterile TSB medium. *L. kalixensis* strain CCUG 48459T was cultured overnight in sterile TSB medium in an anaerobic growth chamber containing Oxoid AnaeroGen (Thermo Fisher Scientific) sachets to create the anaerobic atmosphere. Subsequently, bacteria were diluted 1:100 by adding 2 μL of liquid culture into medium in a sterilized, clear polystyrene 96-well plate to a final assay volume of 200 μL. Subsequently, BzLPRDSar was added to the bacteria from a filter sterilized stock in MQ-H_2_O to give 2, 8, 32 and 128 μg/ml final concentrations. Then, bacteria were incubated at 37 °C for 3, 6, 12, 36 or 48 hours. Plates containing *L. kalixensis* were kept incubating under anaerobic conditions at 5% CO_2_ and 6% O_2_ at 37 °C. Subsequently, their absorbance was measured at 600 nm using a Varioskan Lux microplate reader (Thermo Fisher Scientific). All individual samples were carried out in triplicate.

### Biofilm inhibition assays

For biofilm inhibition experiments *S. aureus* strain CCUG 35571, *E. faecalis* strain CCUG 34289, *B. cereus* strain UW85, *S. agalactiae* strain CCUG 4208T and *S. epidermidis* strain ATCC 35984 were cultured overnight in sterile BHI broth, while *L. kalixensis* strain CCUG 48459T was cultured overnight in sterilized BHI broth under anaerobic conditions in a growth chamber containing Oxoid AnaeroGen sachets. Sterilized, clear polystyrene 96-well plates were prepared by coating with plasma by addition of 20% of final assay volume of Coagulase Test dissolved in 3 mL of sterilized distilled water to the wells and incubating overnight at 4 °C. Then, the cultured strain was diluted 1:100 by adding 2 μL into BHI medium on the plasma coated plate to a total volume of 200 μL. Subsequently, BzLPRDSar was added to the bacteria from filter sterilized stocks in MQ-H_2_O to give 8, 32 and 128 μg/ml final concentrations and incubated for 16 hours at 37 °C without shaking. The *L. kalixensis* strain was kept incubating under anaerobic conditions as described previously. Then, the supernatant was removed, and the wells were rinsed with PBS and 200 μL of 1% crystal violet solution (diluted in PBS) was added and the biofilms were stained for 10 minutes. The wells were washed with PBS twice and 200 μL of a 95% ethanol solution in water was added to solubilize the cells for 30 minutes. 100 μL of suspension was transferred into a new 96-well plate and the absorbance measured at 595 nm using a Varioskan Lux microplate reader (Thermo Fisher Scientific). All individual samples were carried out in triplicate.

### SEM measurements

For SEM imaging, *S. aureus* strain CCUG 3557 and *S. epidermidis* strain ATCC 35984 were cultured overnight in sterile TSB medium. Then, the cultured strain was diluted 1:100 into sterile TSB medium and BzLPRDSar was added to a final concentration of 128 μg/mL. Cells grown in medium without any treatment were used as the control. After 16 hours of incubation, the cells were washed and fixed in 3% glutaraldehyde for 2 hours. Finally, the fixed samples were dehydrated with a series of washes with increasing ethanol concentration (40, 50, 60, 70, 80, 90 and 100%) for 10 minutes each and then dried for 2 hours at room temperature. Before imaging, the dried samples were sputter coated with gold (5 nm). SEM imaging was performed with a Supra 60 VP (Carl Zeiss AG) microscope.

## Supporting information

Hintzen etal_Supplementary Information

## Acknowledgements

The Knut and Alice Wallenberg Foundation via the Wallenberg Centre for Molecular and Translational Medicine (AT), Swedish Research Council (2020-04299 to AT, Cancerfonden (22-2409 to AT) and Centre for Antibiotic Resistance Research (CARe) are gratefully acknowledged. The authors thank Santosh Pandit for providing the multidrug resistant *S. aureus, E. faecalis* and *K. pneumoniae* bacterial strains, Mirjam Dannborg for help with setting up experiments under anaerobic growth conditions and providing the *L. kalixensis* strain.

