## Supplementary material for "Targeted Sortase A Inhibition by Novel Peptidomimetic Antivirulents against Staphylococcal Infections": Hintzen etal_Supplementary Information

### 1. Analytical peptide data:

| Peptide | Calculated Mw [g/mol] | Experimental Mw [g/mol] | ppm | tR [min] |
|---|---|---|---|---|
| BzLPRDSar | 674.3626 | 674.3655 | 4.3 | 5.22 |

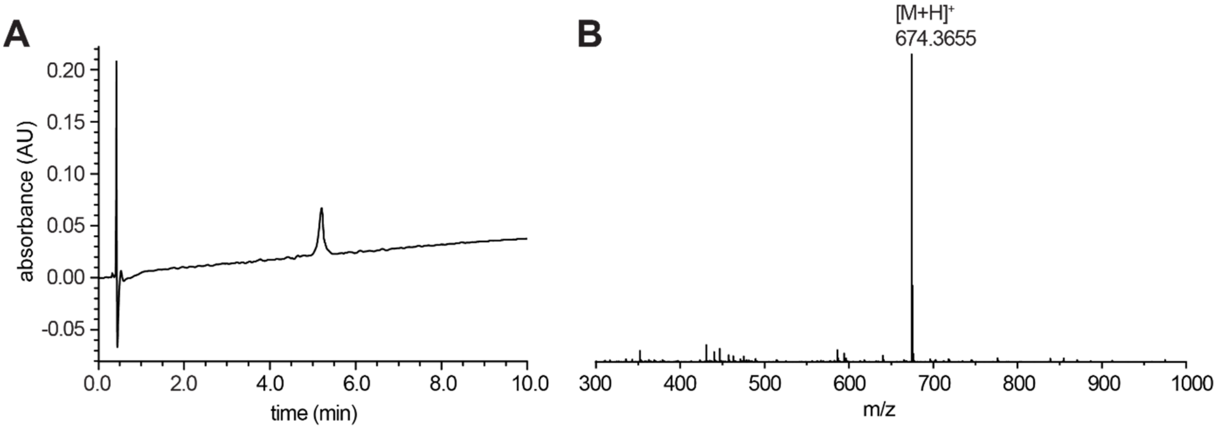


**Fig. S1:** A) HPLC chromatogram (gradient from 5 to 50% ACN in water over 10 min at 2 ml/min, detection at 214 nm) and B) high resolution mass spectrum for peptide 18 BzLPRDSar.

### 2. Growth profiling *K. pneumoniae*:

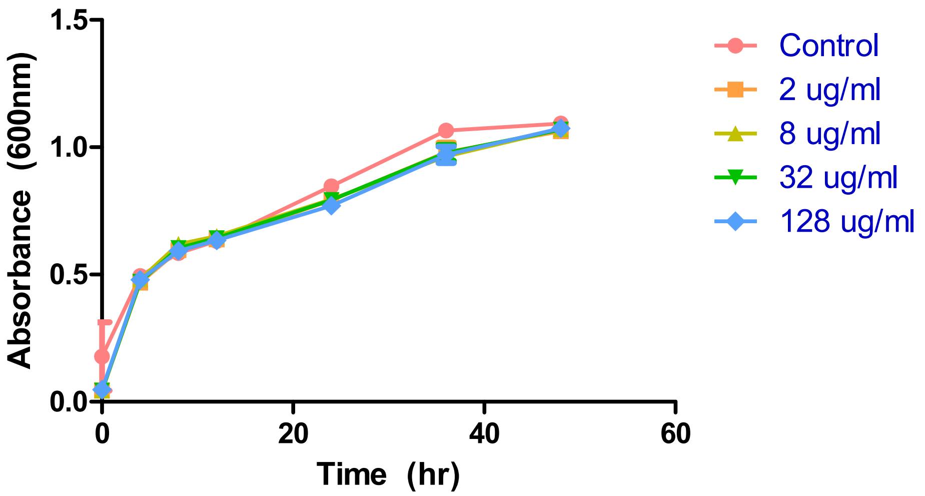


**Fig. S2:** Growth profiling curve of *K. pneumoniae* incubated in presence of varying concentrations of BzLPRDSar. Absorbance was measured at 600 nm as a measure of OD. All samples were measured in triplicate, error bars reported as standard error (±SE).

### 3. Additional SEM images *S. aureus* and *S. epidermidis*:

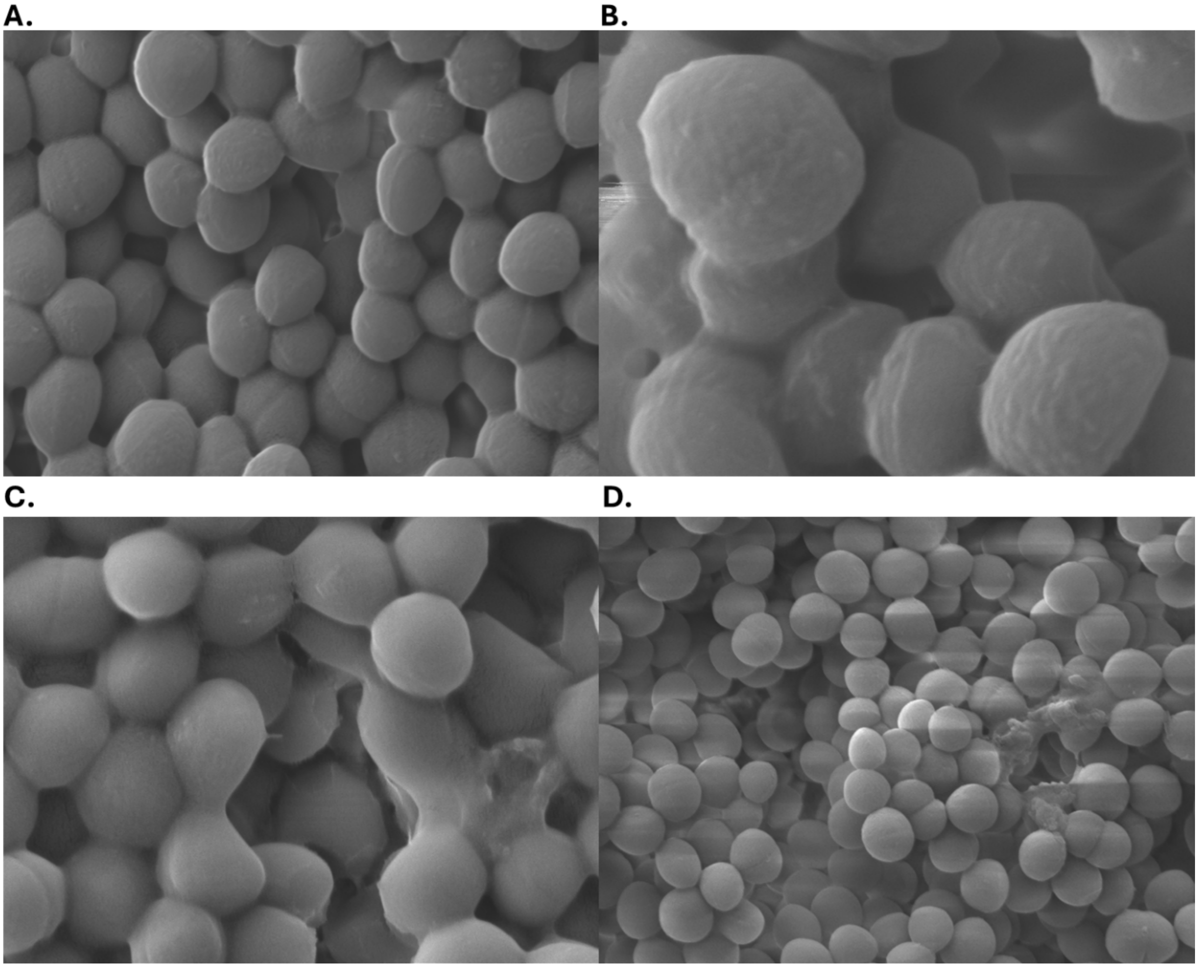


**Fig. S3:** SEM images illustrating structural changes in biofilms of *S. aureus* and *S. epidermidis.* **A)** *S. aureus* biofilms, untreated control, magnification 25,000x; **B)** *S. aureus* biofilms, incubated with 128 µg/ml of BzLPRDSar for 18 hours, magnification 50,000x; **C)** *S. epidermidis* biofilms, untreated control, magnification 25,000x; **B)** *S. epidermidis* biofilms, incubated with 128 µg/ml of BzLPRDSar for 18 hours, magnification 15,000x;
